# Prokaryotic Argonaute from *Archaeoglobus fulgidus* interacts with DNA as a homodimer

**DOI:** 10.1101/2020.05.06.080267

**Authors:** Edvardas Golovinas, Danielis Rutkauskas, Elena Manakova, Marija Jankunec, Arunas Silanskas, Giedrius Sasnauskas, Mindaugas Zaremba

## Abstract

**Background:** Argonaute (Ago) proteins are found in all three domains of life. The best characterized group is eukaryotic Argonautes (eAgos), which are the core of RNA interference. The best understood prokaryotic Ago (pAgo) proteins are full-length pAgos. They are monomeric proteins, all composed of four major structural/functional domains (N, PAZ, MID and PIWI) and thereby closely resemble eAgos. It is believed that full-length pAgos function as prokaryotic antiviral systems, with the PIWI domain performing cleavage of invading nucleic acids. However, the majority of identified pAgos are shorter and catalytically inactive (encode just MID and inactive PIWI domains), thus their action mechanism and function remain unknown.

**Results:** In this work we focus on AfAgo, a short pAgo protein encoded by an archaeon *Archaeoglobus fulgidus*. We find that in all previously solved AfAgo structures, its two monomers form substantial dimerization interfaces involving the C-terminal β-sheets. Led by this finding, we have employed various biochemical and biophysical assays, including single-molecule FRET, SAXS and AFM, to test the possible dimerization of AfAgo. SAXS results confirm that WT AfAgo, but not the dimerization surface mutant AfAgoΔ, forms a homodimer both in the apo-form and when bound to a nucleic acid. Single molecule FRET and AFM studies demonstrate that the dimeric WT AfAgo binds two ends of a linear DNA fragment, forming a relatively stable DNA loop.

**Conclusion:** Our results show that contrary to other characterized Ago proteins, AfAgo is a stable homodimer in solution, which is capable of simultaneous interaction with two DNA molecules. This finding broadens the range of currently known Argonaute-nucleic acid interaction mechanisms.

## INTRODUCTION

Argonaute (Ago) proteins are found in all three domains of life (bacteria, archaea, and eukaryotes). The best characterized group is eukaryotic Ago (eAgo) proteins. Being the functional core of RNA interference machinery, eAgos are involved in regulation of gene expression, silencing of mobile genome elements, and defense against viruses. From the structural and mechanistic point of view, all eAgos are very similar, as they all use small RNA molecules as guides for sequence-specific recognition of RNA targets, and are monomeric proteins sharing four conserved functional domains, which are organized in a bilobed structure [1]. The N-terminal lobe consists of the N-domain that separates guide and target strands [2], and the PAZ domain responsible for binding the 3’-terminus of the guide RNA; the C-terminal lobe consists of the MID domain, which binds the 5’-terminus of the guide RNA, and the PIWI domain, an RNase. Upon recognition of the RNA target, eAgos may either cleave it employing the catalytic activity of the PIWI domain, or, especially eAgo proteins that encode catalytically inactive PIWI domains, recruit partner proteins leading to degradation of the target RNA or repression of its translation [3].

Ago proteins are also identified in 9% of sequenced bacterial and 32% archaeal genomes [4, 5]. Unlike eAgos, which exclusively use RNA guides for recognition of RNA targets, different pAgos may use either RNA or DNA guides and/or targets [6], and may also differ in their structural organization. The best understood prokaryotic Ago (pAgo) proteins are the so called full-length pAgos, which are composed of N, PAZ, MID and PIWI domains, and thus closely resemble eAgo proteins. There is mounting evidence that full-length pAgos function as prokaryotic antiviral systems, with the PIWI domain performing cleavage of invading nucleic acids. However, the majority (~60 %) of identified pAgos are shorter (encode just MID and PIWI domains) and are catalytically inactive due to mutations in the PIWI domain. Though similar artificial truncations of eukaryotic Agos preserve most of functionality characteristic to full-length proteins [7–10], the function and mechanism of the naturally-occurring short catalytically inactive pAgos remains unknown [4, 5].

In this work we focus on the short prokaryotic Argonaute AfAgo encoded by a hyperthermophilic archaeon *Archaeoglobus fulgidus* [4, 5]. Like other short pAgos, AfAgo contains a MID and a catalytically inactive PIWI domains (albeit sequence analysis suggests that AfAgo MID and PIWI domains are closer to those found in full-length, rather than most short pAgos [4, 5]). For over a decade it served as a model system for structural and mechanistic studies of Argonaute-nucleic acids interactions [10–12]. It is also one of the first and the best structurally characterized prokaryotic Argonautes, with an apo- and 3 dsDNA/RNA-bound structures currently available [13–16]. However, its biological role, in part due to lack of catalytic activity, remains elusive. Unexpectedly, inspection of these structures revealed that regardless of the crystal form and symmetry, two AfAgo subunits always form a substantial dimerization interface involving N-terminal residues and/or β-strands located close to the C-termini. Using various biochemical and biophysical assays, including single-molecule FRET, small angle X-ray scattering (SAXS), and atomic force microscopy (AFM), we show that AfAgo is indeed a stable homodimer in solution, and is capable of simultaneous interaction with two DNA molecules. This broadens the range of currently known interaction mechanisms involving nucleic acids and Argonaute proteins.

## RESULTS

### AfAgo is a homodimer in the available X-ray structures

AfAgo is a 427 amino acid (aa) 49.2 kDa prokaryotic Argonaute protein found in the hyperthermophilic archaeon *Archaeoglobus fulgidus*. To date, four AfAgo structures, both of the apo-form and bound to DNA and RNA duplexes were solved [13–16]. AfAgo monomer is composed of two major domains, the N-terminal MID (residues 38-168), and the C-terminal PIWI (residues 168-427) [13]. The MID domain specifically binds the 5’-phosphorylated end of the presumed guide DNA/RNA strand, and also makes contacts to the complementary target DNA/RNA strand [14–16]. The PIWI domain makes contacts to both guide and target DNA/RNA strands, but is catalytically inactive due to mutations in the RNase H-like catalytic center. Unexpectedly, inspection of available AfAgo structures (Supplementary table S4) revealed that in all structures known so far, AfAgo subunits form homodimers. The dimerization interface in the AfAgo-dsRNA structure (PDB ID 1ytu) is asymmetric, and primarily involves the C-terminal β-strands (residues 296-303) from both subunits present in the asymmetric unit that together form a parallel β-sheet, and the N-terminal residues from one of the subunits (Figure 1A). The dimer formed in this case is compact (henceforth, a ‘closed’ dimer). In contrast, dimerization interfaces in three other cases (PDB IDs 1w9h, 2bgg and 2w42) are nearly symmetrical with respect to the secondary structure elements involved (albeit in PDB IDs 2bgg and 2w42 they belong to different protein chains present in the asymmetric unit): the C-terminal β-strands form 8-strand β-barrels, with the sheets from different subunits interacting via strands β14 (residues 297-302) and β15 (residues 314-318, Figure 1B). The resultant dimers are less compact (henceforth, ‘open’ dimers).

**Figure 1.**
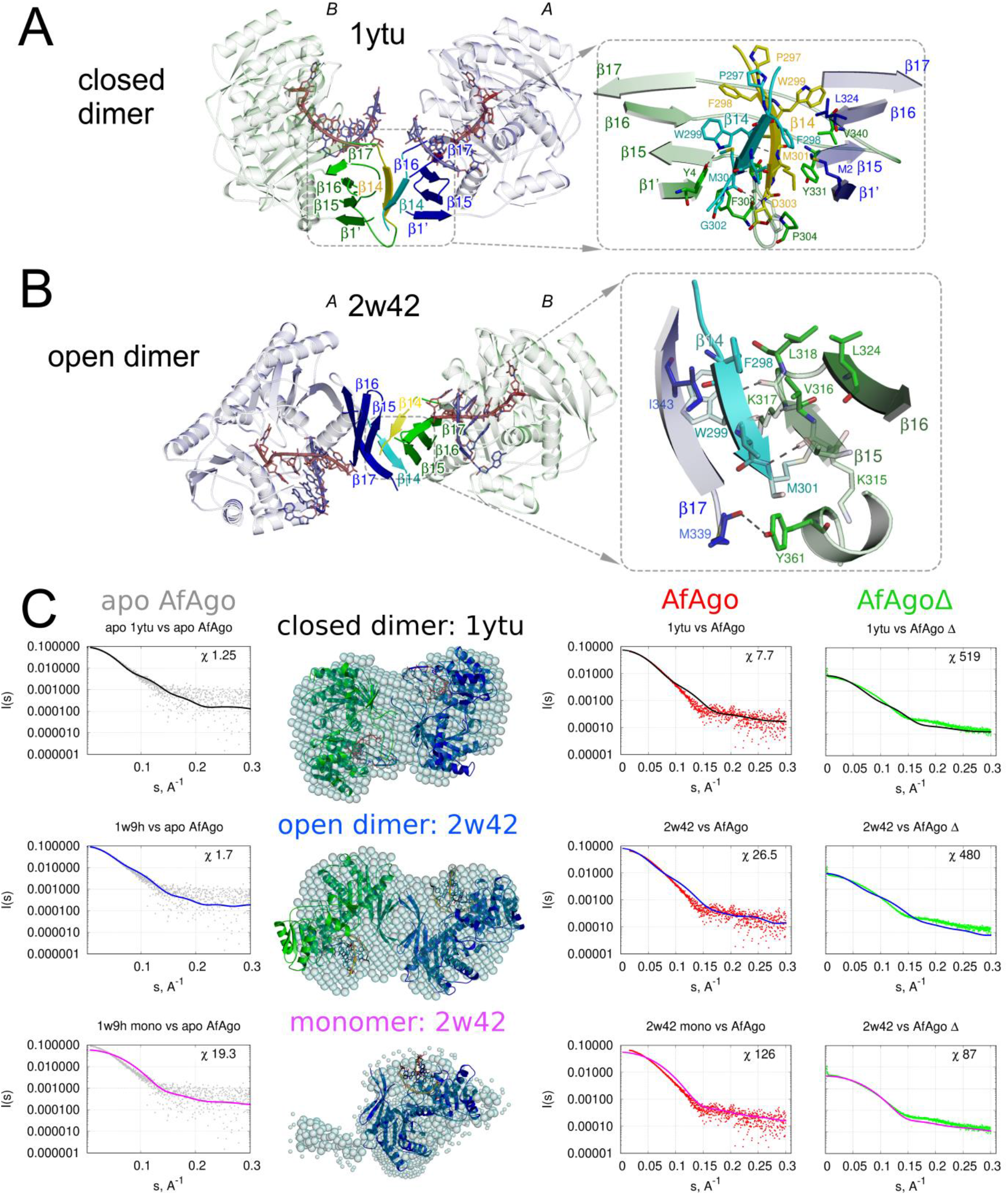
Dimerization of AfAgo. (A-B). Protein subunits are colored blue (protein chain *A*) and green (protein chain *B*). The interface-forming secondary structure elements are highlighted and numbered according to the PDB ID 2w42 assignment made by PDBsum [50]. The ‘guide’ DNA/RNA strands bound by AfAgo are colored red, ‘target’ strands — blue. Residues 296-303 deleted in AfAgoΔ are colored cyan and yellow. Hydrogen bonds are shown as dashed lines. (A) AfAgo complex with dsRNA (PDB ID 1ytu, both protein chains as present in ASU), the ‘closed’ dimer [14]. (B), AfAgo complex with dsDNA (PDB ID 2w42, protein chain B is produced by operator (X,Y,Z+(−1 2 2)&{0 −1 - 1}) [16]) - the ‘open’ dimer. β-strands from both subunits assemble into a closed β-barrel structure, with intersubunit interface formed by β14 and β15 strands of neighboring subunits. (C) SAXS data of WT AfAgo apo and complex with MZ-1289 DNA (grey and red points, respectively), and the dimerization mutant AfAgoΔ with MZ-1289 DNA (green points, right column) are compared with the scattering curves generated from the ‘closed’ dimer with or without dsRNA (PDB ID: 1ytu, black curves), ‘open’ dimer (PDB ID: 2w42 or 1w9h for apo AfAgo, blue curves) and monomeric apo and AfAgo-DNA complex (PDB ID: 2w42 or 1w9h, magenta curves) by CRYSOL. Corresponding AfAgo structures are shown in the second column superimposed with the dummy atom models generated using the SAXS data of AfAgo complex with MZ-1289 oligoduplex.

The solvent accessible surface areas buried at the dimerization interfaces, as calculated by the PISA server (https://www.ebi.ac.uk/pdbe/pisa/pistart.html, [17]) and the number of inter-subunit H-bonds (Supplementary table S4) in both ‘open’ and ‘closed’ dimers are typical for stable protein dimerization interfaces [18, 19]. This observation prompted us to test the oligomeric state, the possible dimerization mode, and mechanism of nucleic acid binding of AfAgo in solution using various biochemical and biophysical techniques. For that purpose, we used two variants of AfAgo: the full-length wild-type protein (henceforth, WT AfAgo), and a dimerization mutant AfAgo lacking the 296-303 amino acid residues involved in dimerization (henceforth, AfAgoΔ). Both proteins were successfully purified as described in Materials and Methods, albeit the AfAgoΔ variant was prone to aggregation. Stability of both proteins was considerably improved upon addition of a phosphorylated blunt-end DNA oligoduplex MZ-1289 (Supplementary table S1).

### SAXS measurements

To determine the solution conformation and oligomeric state of AfAgo, we have performed small angle X-ray scattering (SAXS) measurements using DNA-bound and DNA-free full-length WT AfAgo protein and the DNA-bound dimerization mutant AfAgoΔ. Two types of data analysis were performed: (i) the *ab initio* shapes of the proteins in solution were calculated and superimposed with the X-ray AfAgo structures, and (ii) the theoretical scattering data was calculated for the crystallized AfAgo monomer, ‘open’ (PDB ID: 2w42 and 1w9h) and ‘closed’ (PDB ID: 1ytu (with and without bound dsRNA)) dimers, and compared to experimental SAXS scattering data of AfAgo and AfAgoΔ (Figure 1 **Error! Reference source not found**.). The ‘closed’ AfAgo dimer fits the DNA-bound and DNA-free WT AfAgo SAXS data better than the ‘open’ dimer, as judged from the real space fit and the **χ**^2^ (Figure 1C) parameters that reflect the agreement between scattering functions of corresponding crystal structures and SAXS experiments (Figure 1C). As expected, AfAgo monomer gave the best fit to the AfAgoΔ SAXS data (Figure 1C, right column). The SAXS molecular weights calculated for AfAgo (between 94.2 and 106.9 kDa (108-133 kDa for apo AfAgo, that was prone to aggregation), Supplementary table S3) agreed with the expected mass of the dimer complexed with dsDNA (110.5 kDa). The SAXS MW for the AfAgoΔ (between 55.4 and 67.9 kDa) confirmed the monomeric state of the dimerization mutant-dsDNA complex.

### Direct visualization of AfAgo-induced DNA loops by AFM

AfAgo and DNA were deposited on APS-mica and imaged using tapping AFM. A typical AFM image of AfAgo-DNA complexes is shown in Figure 2. Several types of protein-DNA complexes, shown as enlarged insets in Figure 2, were observed: (i) linear DNA with a protein molecule bound to one DNA end; (ii) linear DNA with protein molecules bound to both DNA ends; (iii) ring-shaped (looped) DNA. Other species, including naked DNA, or more complex structures, involving, e.g., protein bound to two DNA fragments, were also observed, but were not quantified. We find that the relative distribution of different complexes varied dramatically for WT AfAgo and the dimerization mutant AfAgoΔ (Table 1). The ring-shaped DNA-protein complexes are the dominant species observed with WT AfAgo (55% or 114 out of 208 complexes). The minor fraction of DNA molecules had either protein bound to one end (29%, 61 out of 208 complexes) or to both ends (16%, 33 out of 208). In the case of AfAgoΔ, the majority of complexes had protein bound to both DNA ends (59%, or 187 out of 319, Table 1), and a much lesser fraction (12%, or 38 out of 319) were ring-shaped structures. Since AfAgoΔ lacks the dimerization interface observed in the structures, the observed small fraction of ring-shaped DNA complexes could be due to sample treatment with glutaraldehyde (see Materials and Methods for details), which could occasionally lead to inadvertent cross-linking of two AfAgoΔ monomers bound to termini of the same DNA molecule, thereby generating looped complexes.

**Figure 2.**
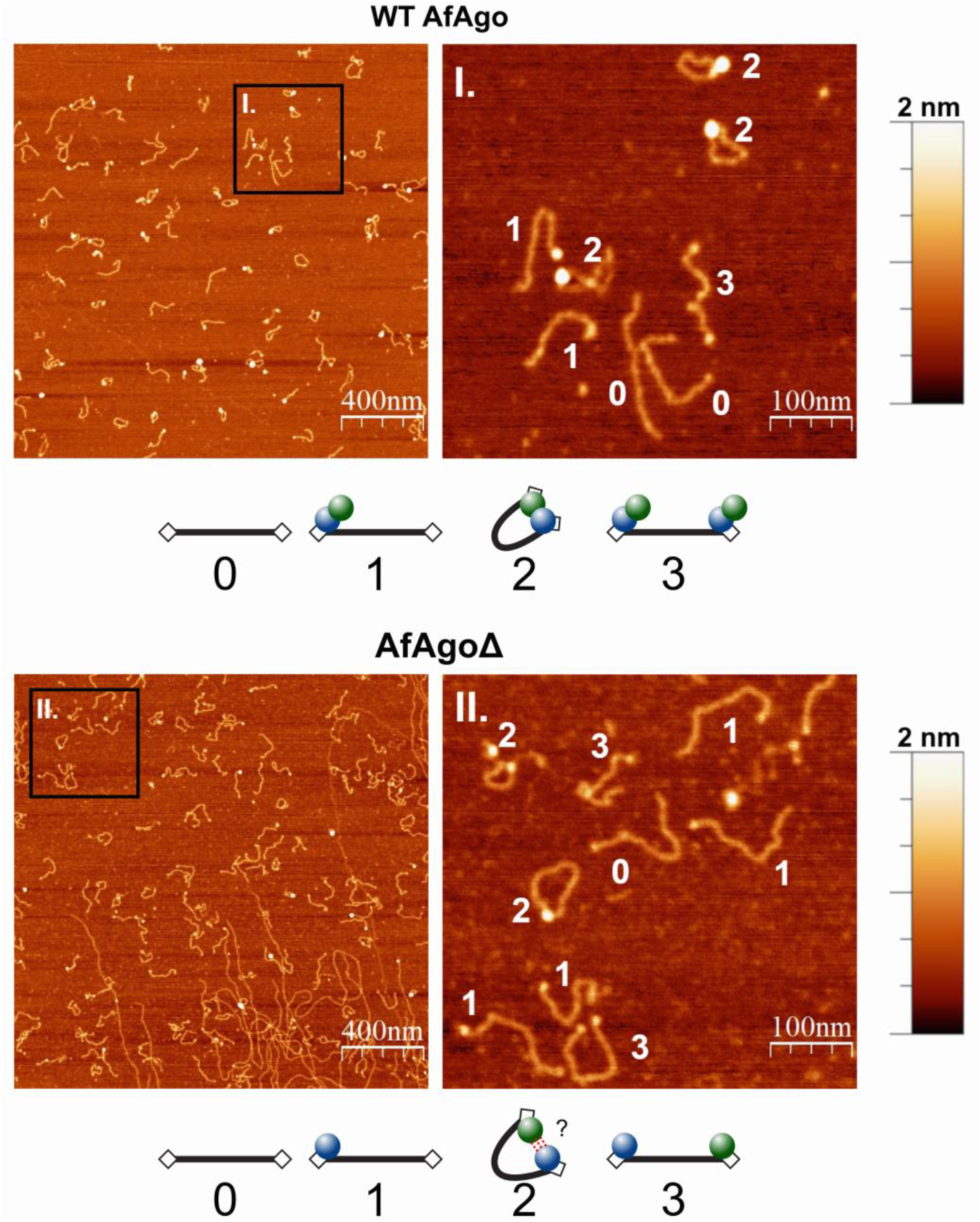
Visualization of AfAgo-induced DNA loops by AFM. Representative AFM images and enlarged views of DNA-protein complexes adsorbed to APS-mica acquired in the air are shown. DNA+WT AfAgo (top) and DNA + AfAgoΔ (bottom), area of each left column image is 4 μm^2^, scale bar is 400 nm. Right column shows regions marked by squares in the respective images on the left, enlarged 4-fold, scale bar 100 nm. Various observed species are depicted below the respective images and are marked as follows: ‘0’ - naked DNA; ‘1’ - protein bound to one DNA end; ‘3’ - protein bound to both DNA ends; ‘2’ - ring-shaped (looped) DNA. In the case of AfAgoΔ, species ‘2’ is presumably formed due to glutaraldehyde crosslinking. The Z scale bar is 2 nm.

**Table 1.** AfAgo-DNA complexes observed by AFM.

| Protein | DNA loops, % | Linear |  | Total number of analyzed complexes |
| --- | --- | --- | --- | --- |
|  |  | *Bound to one end, %* | *Bound to both ends, %* |  |
| WT AfAgo | 55 | 29 | 16 | 208 |
| AfAgoΔ | 12 | 29 | 59 | 319 |

### WT AfAgo induces DNA loops in solution

Dimerization of WT AfAgo observed in X-ray structures and SAXS measurements is unique among Argonaute proteins. It raises a question if the AfAgo/DNA stoichiometry observed in the structures, an AfAgo dimer bound to two nucleic acid molecules, is also relevant in solution. To address this question we have examined the interaction of AfAgo with a DNA substrate bearing two binding sites exploiting the method of single-molecule Förster resonance energy transfer (smFRET). If AfAgo binds DNA as a dimer, as observed in the X-ray structures, it should be capable of interacting with both DNA ends simultaneously, inducing a DNA loop, which, in addition to AFM, could be monitored as a change of FRET efficiency between dyes tethered close to DNA ends (Figure 3A). Utilization of a single dual-labeled two-target DNA substrate (rather than two short DNA duplexes carrying different fluorescent labels) increases the probability of AfAgo interaction with both DNA targets at low reactant concentrations required for the single-molecule setup.

**Figure 3.**
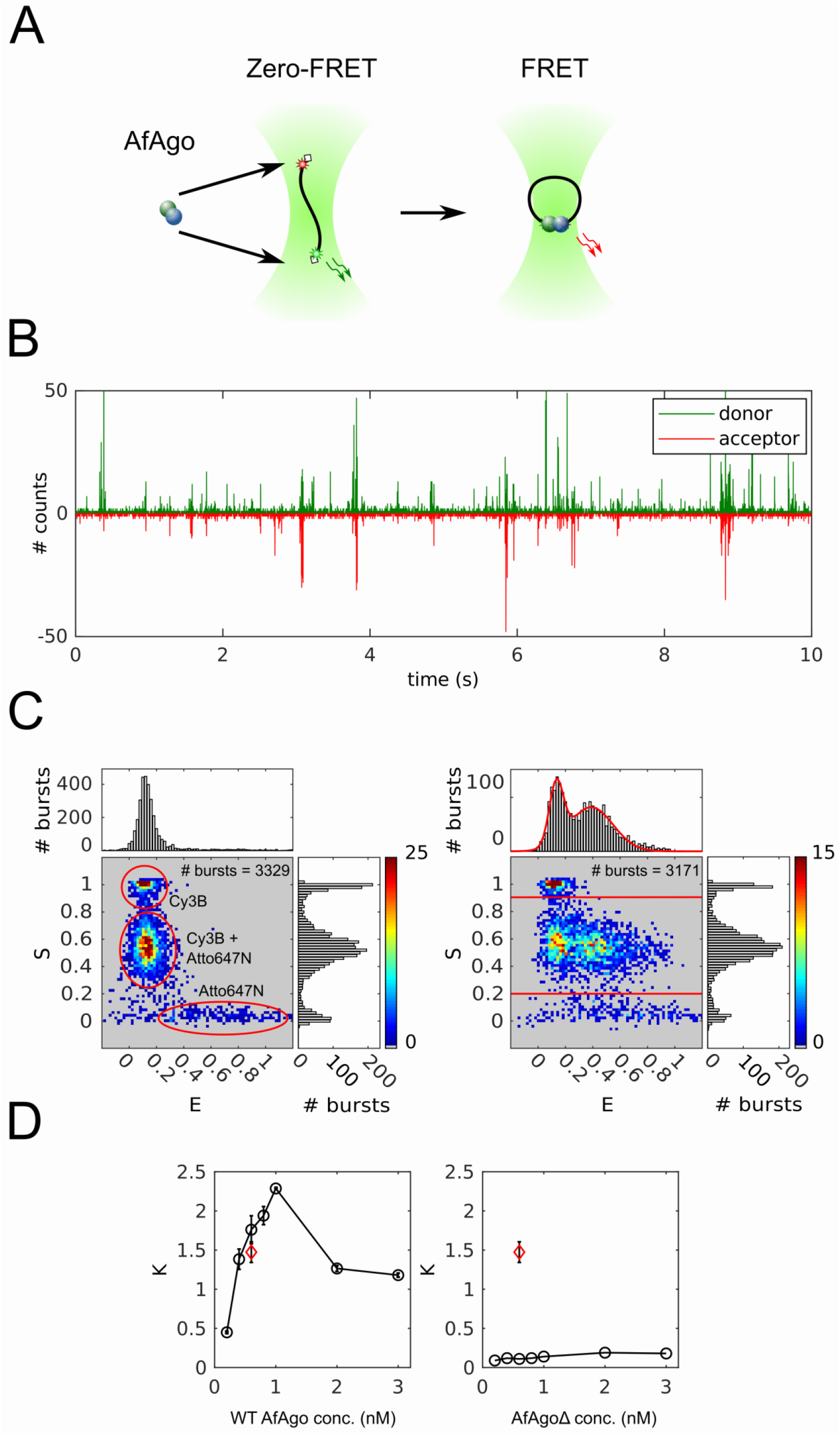
Single-molecule studies of AfAgo-DNA interactions in solution. (A) A schematic overview of the single molecule assay. Left, free DNA, right, DNA-WT AfAgo (blue and green circles) complex in a looped state. (B) Fluorescence intensity trace with 1 ms time bin of 25 pM DNA with 1 nM AfAgo. Donor fluorescence upon donor excitation is presented in the upper part of the graph. Inverted acceptor fluorescence upon donor excitation is presented in the lower part of the graph. (C) Left - E-S histogram of DNA alone. The top and side axes contain, respectively, one-dimensional E (proximity ratio) and S (donor/acceptor stoichiometry) histograms of all bursts. Denoted are areas corresponding donor-only DNA, acceptor-only DNA, and dual-labeled DNA. Right - E-S histogram of DNA with 1 nM AfAgo. The one-dimensional E histogram on top is derived from bursts with S = 0.2-0.9, designated by horizontal lines in the E-S histogram. The red curve is a two-Gaussian fit to the data that gave the positions of the Gaussian maxima on the E-axis (0.13±0.01 and 0.39±0.02). (D) Left - dependence of the ratio of looped and unlooped DNA molecules (parameter K) on WT AfAgo concentration (open circles). Right - the dependence of K on the AfAgoΔ concentration (open circles). The red diamonds in both graphs represent the competition experiment performed with 0.6 nM WT AfAgo dimer and 0.6 nM AfAgoΔ monomer. All data points are average values of three measurements ±1 standard deviation.

We have designed a 569 bp DNA construct, which was labelled with a pair of FRET fluorophores, Cy3B and Atto647N, each attached to thymine bases 3 nt away from the respective DNA termini via a C6 linker (Supplementary figure S2). The positions of FRET labels were selected such that upon binding of both DNA ends by an AfAgo dimer, the distance between the label attachment sites (irrespective of the AfAgo dimerization mode), is favorable for FRET (Supplementary figure S4). To promote tighter binding, both DNA ends were phosphorylated, and the 5’-terminal nucleotides were adenines, since AfAgo, like Argonaute CbAgo from *Clostridium butyricum* [20], has a preference for a 5’-terminal A (publication in preparation).

AfAgo interaction with the DNA fragment was monitored by analyzing the fluorescence bursts of single diffusing DNA fragments (Figure 3B). For each DNA molecule we have calculated two parameters. The first parameter S represents the stoichiometry of different fluorophores present on the DNA, and is equal to the ratio I_d_/(I_d_ + I_a_^a^), where I_d_ is the total donor and acceptor intensity upon donor excitation, and I_a_^a^ is acceptor intensity upon acceptor excitation. The relative excitation intensities of the donor and acceptor fluorophores were adjusted such that the stoichiometry parameter S was about 0.5 for DNA molecules labeled with both fluorophores, approx. 0 for the acceptor-only DNA, and close to 1.0 for the donor-only DNA. The second parameter, the proximity ratio E, is equal to I_d_^a^/(I_d_^a^ + I_d_^d^), where I_d_^a^ and I_d_^d^ are acceptor and donor intensities upon donor excitation, respectively. It is expected to be higher for looped DNA molecules with the ends brought into close proximity than for unlooped DNA molecules.

The E-S histogram of DNA alone (Figure 3C, left) exhibits a prominent population with low E and intermediate S values, which corresponds to dual-labeled unlooped (zero-FRET) DNA molecules. The two minor populations observed in the histogram correspond to donor-only (low E/high S) and acceptor-only (high E/low S) DNA fragments. The E-S histogram of DNA in the presence of WT AfAgo exhibits an additional population (intermediate S and intermediate E, Figure 3C, right), which presumably represents DNA molecules looped by WT AfAgo. For further analysis we chose the dual-labeled DNA molecules (S between 0.2 and 0.9) and built their E histogram. The histogram exhibits peaks of near-zero and high E, corresponding to unlooped and looped DNA molecules, respectively (Figure 3C, right).The fit of this histogram with a sum of two Gaussian functions allowed us to calculate the ratio K defined as the ratio of the fraction of looped DNA molecules (the area under the Gaussian with the high E center) and unlooped DNA molecules (the area under the Gaussian with a near-zero E center) over the population of the dual-labeled DNA fragments.

We have measured the ratio K at different WT AfAgo concentrations (Figure 3D). It increased monotonously with increasing WT AfAgo concentration until it reached the maximum value of 2.5 (corresponds to about 70% of looped DNA molecules) at 1 nM WT, but decreased as the protein concentration was increased further.

A similar set of single molecule experiments was performed with the dimerization mutant AfAgoΔ. As shown in Figure 3D, the ratio K at all AfAgoΔ concentrations tested was close to zero, indicating that AfAgoΔ was unable to induce DNA loops. Lack of DNA looping was not due to lack of DNA binding, as shown by electrophoretic mobility shift assay (Supplementary figure S5). Moreover, AfAgoΔ competes with WT AfAgo for DNA ends, as the K value observed in a competition experiment performed with equal concentrations of WT AfAgo dimer and AfAgoΔ monomer was considerably lower than in an experiment with WT AfAgo alone (Figure 3D). Taken together, our results indicate that WT AfAgo dimer is capable of simultaneous interaction with two DNA ends in solution.

### Dynamics of WT AfAgo-induced DNA loops

To explore the dynamics of the WT AfAgo-induced DNA looping events, we have used total internal reflection fluorescence (TIRF) microscopy to perform single molecule FRET experiments on surface-immobilized DNA (Figure 4). For that purpose we used a DNA fragment that was essentially identical to the one used for single-molecule studies in solution, except that it carried a biotin 386 bp away from the donor end for surface immobilization (Supplementary figure S2). After verifying that WT AfAgo induces loops on this substrate in solution (Supplementary figure S6B), we immobilized the biotinylated DNA on a surface and then recorded fluorescence movies in the absence or in the presence of WT AfAgo (Supplementary figure S6A). From each frame of the movie we then extracted donor and acceptor intensities for individual DNA fragments, selected trajectories with anti-correlated changes of the donor and acceptor intensities (indicating the occurrence of FRET) and calculated the time courses of the proximity ratio, E. An example of such a trajectory is presented in Figure 4C. In a control with no AfAgo we could find no DNA fragments exhibiting FRET (Figure 4B).

**Figure 4.**
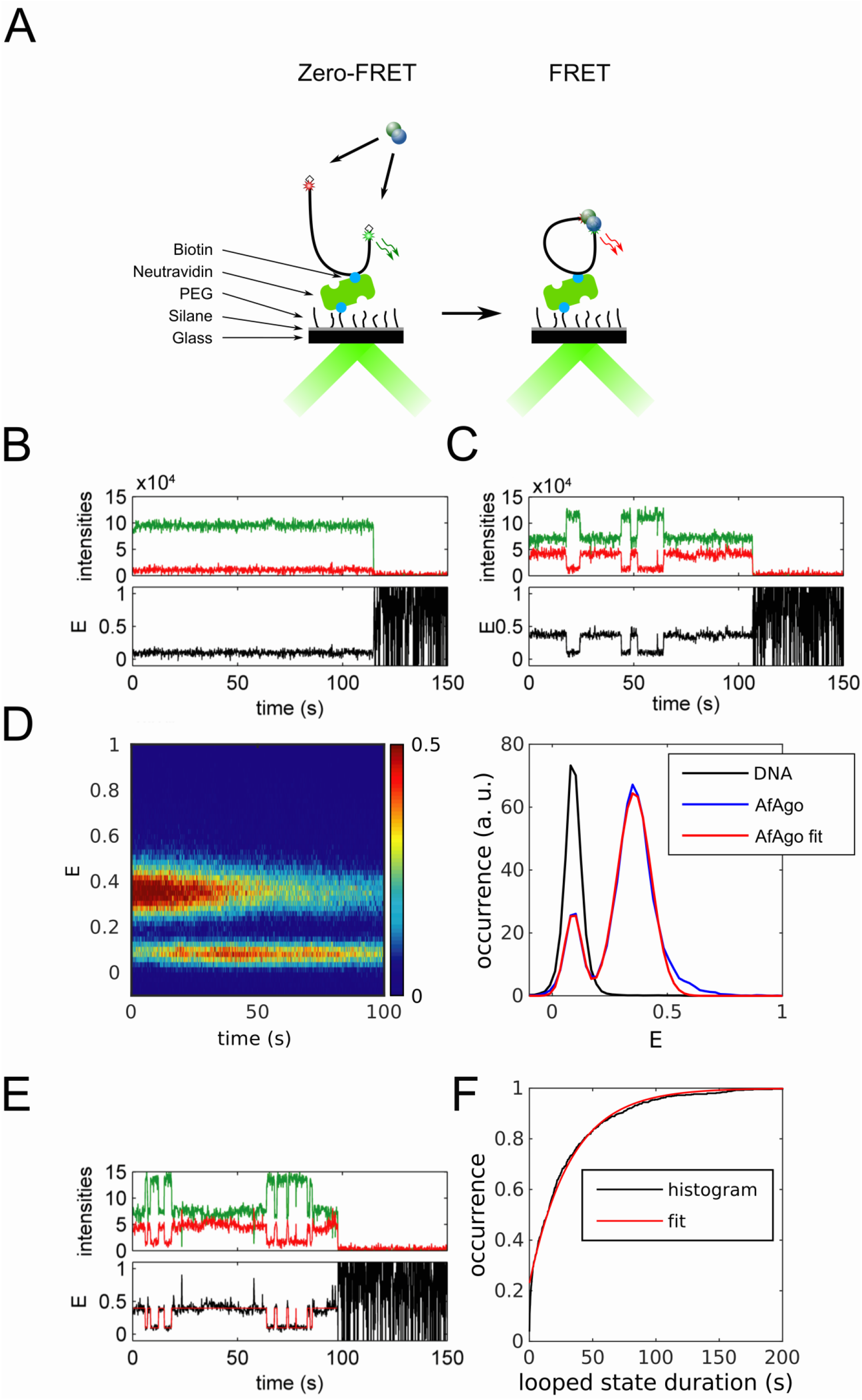
Dynamics of WT AfAgo-induced DNA loops. (A) A schematic overview of the single-molecule assay using TIRF microscopy. (B, C) Trajectories of donor (green) and acceptor (red) intensity and corresponding proximity ratio, E, of individual DNA fragments without (B) and with 10 nM AfAgo (C). (D) Left - an image of 287 pooled time traces of the proximity ratio, E, from the measurement with 10 nM of AfAgo. The image is normalized to the maximum image intensity. Right – a section of the image in the left integrated over the first 10 s shown with the two-Gaussian fit. The positions of the Gaussian maxima are 0.09±0.01 and 0.36±0.01. For comparison a trace- and time-averaged section from the measurement of 227 traces on bare DNA is shown. (E) An example of trajectories of donor (green) and acceptor (red) intensity and corresponding proximity ratio, E, with HMM idealization of an individual DNA fragment with 10 nM AfAgo. F – cumulative histogram of the looped state durations from 287 E traces with a single-exponential fit with the exponential factor of 33±1 s.

To assess the average values of different E levels apparent in individual trajectories we pooled the selected E trajectories and averaged the resulting image within the first 10 s (Figure 4D, left). The single-molecule population and time-averaged E exhibits two peaks with maxima at 0.09 and 0.36, corresponding to the unlooped and looped DNA molecules, respectively (Figure 4D, right). These E values are also in good agreement with the E values obtained from the measurement in solution (Figure 3C).

A superficial inspection of E trajectories of individual DNA fragments revealed that their looping dynamics are rather diverse. There exist trajectories with the looped state lasting the whole measurement, whereas other trajectories are more dynamic (Figure 4C, Supplementary figure S6C, D), exhibiting a number of transitions between the looped and unlooped states. The looped E state also exhibits more subtle dynamics (Supplementary figure S6D) which could probably be attributed to the conformational flexibility of AfAgo at the dimerization interface.

To quantify the looped state duration we first idealized the E trajectories using HMM in QuB software (Figure 4E). Then, from the idealized trajectories we built the cumulative histogram of the looped state durations (Figure 4F). The trajectory edge dwells were not omitted in order to preserve the information on the occurrence of states lasting during the whole trajectory. The exponential factor of a single-exponential fit of the cumulative histogram was 33±1 s. The maximum recorded looped state duration is, however, limited by the duration of our measurement (200 s) and the duration of the fluorescent state of the fluorophores before photobleaching. The value of the exponential factor thus sets the lower limit of the looped state duration.

## DISCUSSION

All characterized long Argonaute proteins interact with their RNA and/or DNA targets as monomers, binding a single copy of each guide and target nucleic acids. Surprisingly, we reveal here that AfAgo, a prokaryotic Argonaute from the hyperthermophilic archaeon *Archaeoglobus fulgidus*, follows a different mechanism, which involves homodimerization and simultaneous interaction with two guide-target nucleic acid duplexes.

First, we show that AfAgo is a homodimer in all previously solved X-ray structures, including apo-protein, and complexes with RNA and DNA (Supplementary table S4). Two types of AfAgo dimerization interfaces formed by the C-terminal β-sheets are observed in the structures. Both types of interfaces bury a comparable solvent-accessible surface area (Supplementary table S4), but result in distinct arrangement of AfAgo subunits relative to one another, which we term ‘closed’ and ‘open’ dimers (Figure 1 A and B, respectively). The ‘closed’ type of AfAgo homodimer, formed when the interface involves both the N-terminal residues and the C-terminal β-strands (Figure 1A), provides a better fit to our SAXS data, suggesting that it is the major type of DNA-bound WT AfAgo dimer present in solution (Figure 1C). It remains to be determined if formation of the alternative ‘open’ dimer observed in several structures (Figure 1B and Supplementary table S4) was influenced by crystal packing, or rather it is an alternative less abundant arrangement of AfAgo subunits that co-exists in solution at equilibrium with the ‘closed’ form. As expected, removal of the β-strands located at the intersubunit interface (variant AfAgoΔ) renders AfAgo incapable of dimerization (Figure 1).

Simultaneous binding of WT AfAgo homodimer to both ends of a linear DNA fragment would result in a DNA loop. Formation of such looped DNA molecules upon incubation with WT AfAgo was directly visualized using AFM (Figure 2); as shown in Table 1, they constitute the majority of all protein-DNA complexes detected. As expected, very few looped DNA molecules were observed in similar experiments performed with the dimerization mutant AfAgoΔ (Table 1). Thus, AFM provides further proof that homodimeric WT AfAgo is capable of simultaneous interaction with two DNA ends, while the dimerization mutant AfAgoΔ, being a monomer, binds just one DNA end.

To further characterize the interaction of WT AfAgo with DNA in solution, we have performed single-molecule FRET measurements (Figure 3) using a 569 bp DNA fragment labelled with different fluorescent labels (a FRET pair) close to DNA ends. Design of the DNA substrate ensured that binding of WT AfAgo dimer to both DNA ends would bring the fluorophores into close proximity, resulting in FRET. Comparison of donor/acceptor channel records for free DNA and DNA with either WT AfAgo or dimerization-incapable AfAgoΔ confirmed that WT AfAgo forms DNA loops, and that loop formation requires AfAgo dimerization.

At least two mechanisms can be proposed for the formation of the WT AfAgo dimer / looped DNA complex. The first mechanism is valid if apo-WT AfAgo forms a stable dimer. The reaction (Figure 5) proceeds via (i) association of free DNA (species ‘0’) with a single WT AfAgo dimer, which binds to one DNA end (species ‘1’); (ii) capture of the second DNA terminus by the DNA-bound AfAgo in an intramolecular reaction, resulting in a looped complex (species ‘2’); (iii) alternatively, association of the second WT AfAgo dimer with the unoccupied target (the second DNA end) of species ‘1’ leads to species ‘3’, which is no longer capable of loop formation. Such mechanism was demonstrated for many proteins capable of DNA looping, including restriction endonucleases [21–23] and transposases [24–27]. The second mechanism is valid if apo-WT AfAgo is a monomer in solution, and dimerizes only upon binding to DNA. In such a case the following steps lead to a looped complex: (i) a single AfAgo monomer binds the first DNA end (species ‘4’, Figure 5); (ii) the second AfAgo monomer binds the second end (species ‘5’); (iii) two DNA-bound monomers associate forming the looped complex ‘2’ (Figure 5). Such mechanism is less common, albeit it was also demonstrated for some nucleic acid enzymes [28]. Single molecule FRET experiments in solution allowed us to distinguish the two above mechanisms. We have found that the amount of the looped (high-FRET) complex is dependent on AfAgo concentration. It increases until an optimal protein concentration is reached (approx. 1 nM dimer in our experimental setup, Figure 3D), and steeply declines upon further increase in WT AfAgo concentration. Such behavior is expected only for the mechanism involving homodimeric WT AfAgo (species ‘0’, ‘1 ‘, ‘2’, and ‘3’, Figure 5), as excessive protein concentrations favor binding of two WT AfAgo dimers to both DNA ends (species ‘3’), and thereby hinder loop formation. No such loop disruption at increased protein concentrations is expected for the second mechanism involving monomeric apo-WT AfAgo (species ‘0’, ‘4’, ‘5’, and ‘2’), as the species favored at the highest protein concentrations (‘5’) is still capable of loop formation. Taken together, our single-molecule studies in solution favor the DNA looping mechanism involving a stable apo-WT AfAgo dimer (species ‘0’, ‘1 ‘, ‘2’, ‘3’, Figure 5), a finding that is also consistent with SAXS measurements of apo-WT AfAgo (Table 1, Figure 1C). However, we cannot completely rule out mixed mechanisms involving an equilibrium of monomeric and dimeric apo-WT AfAgo forms and DNA simultaneously bound to both monomeric and dimeric AfAgo proteins (species ‘6’, Figure 5).

**Figure 5.**
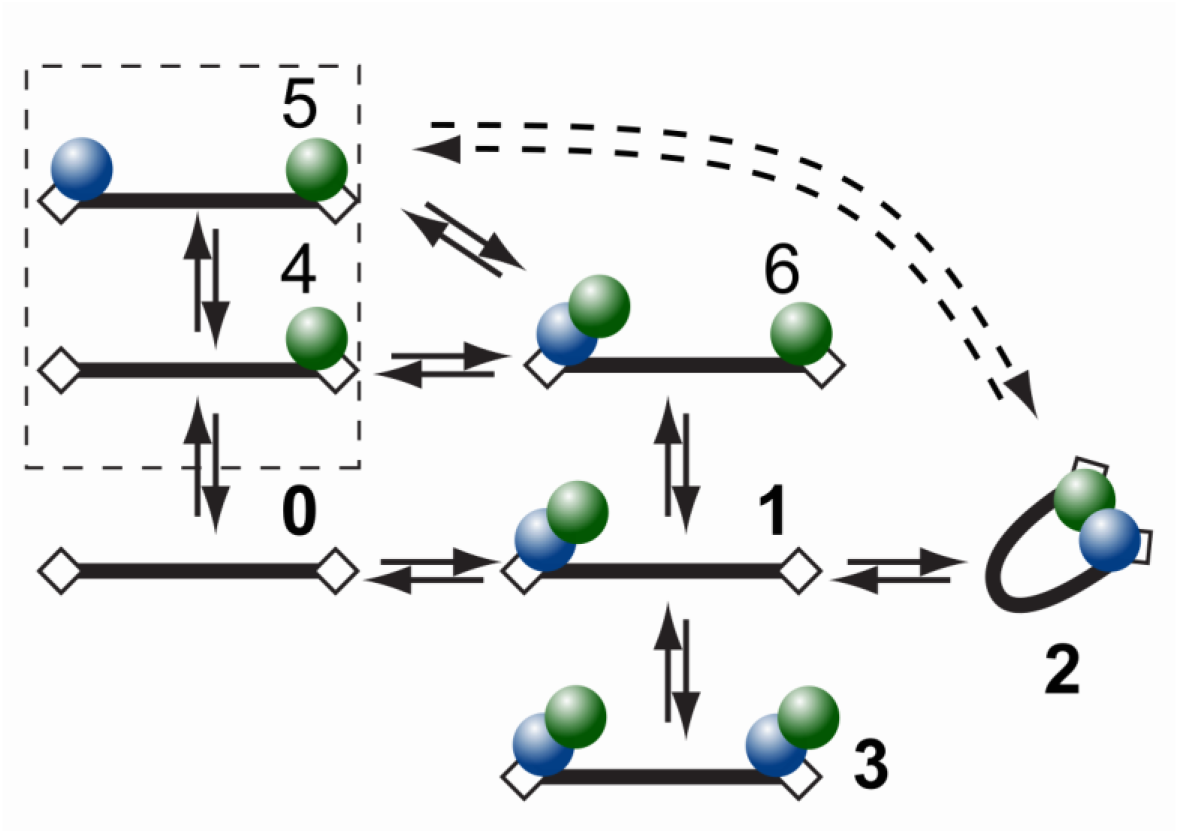
Kinetic schemes depicting possible reaction pathways between two-target site DNA fragment and AfAgo. Black bars represent DNA, rectangular boxes – AfAgo-binding targets (DNA ends), green and blue circles – AfAgo monomers. The 0-1-2-3 pathway represents the case of a stable AfAgo dimer and is supported by our experimental data. Pathway 0-4-5-2 represents the reactions that would occur if AfAgo was a monomer assembling into a dimer upon DNA binding. Dashed box represents species limited to AfAgoΔ.

Single-molecule measurements on immobilized DNA allowed us to assess the dynamic properties of WT AfAgo-induced DNA loops. We find that (i) the DNA loops induced by WT AfAgo are relatively stable, with the lower limit estimate for the loop duration exceeding 30 s (Supplementary figure S6C); (ii) the proximity ratio E of the looped complexes changes over time, suggesting intrinsic dynamics of the AfAgo dimer, presumably attributable to the flexible dimerization interface (Supplementary figure S6D).

## CONCLUSIONS

The ability of WT AfAgo to form homodimers and bring two nucleic acid fragments into close proximity, to the best of our knowledge, is unique among Argonaute proteins, and raises additional questions regarding the currently unknown AfAgo function. Simultaneous interaction with two target sites in the case of restriction endonucleases is believed to increase specificity by preventing inadvertent cleavage of lone unmodified target sites [23, 29]. However, since AfAgo has no intrinsic nuclease activity, it cannot be directly involved in host defense against invading nucleic acids, as recently proposed for the catalytically active full-length pAgos [30, 31]. Instead, the ability of WT AfAgo to form stable synaptic complexes with two DNA ends is reminiscent of transposases [24–27], Cas1-Cas2 integrases [32, 33], and (retro)viral integrases [34, 35], which often bring the reactive 3’-OH groups of two DNA ends into proximity of the integration site. AfAgo thus could serve as the recognition module for the integrated DNA fragment; target DNA recognition, binding and catalysis of the integration reactions would require involvement of additional, currently unknown, partner proteins. In order to test this hypothesis, we currently perform structural and functional studies of other AfAgo-like short prokaryotic Argonautes, and also try to identify and characterize their putative partners.

## METHODS

### Protein expression

The gene encoding WT AfAgo was amplified from *Archaeoglobus fulgidus* genomic DNA by PCR and cloned into a pETDuet vector, yielding a construct with an N-terminal (His)_6_ tag (N-terminal protein sequence MGSSHHHHHHSQDP followed by 1-427 aa of WT AfAgo sequence). The deletion in the dimerization mutant AfAgoΔ was constructed via overlap extension PCR by using two primer pairs, MZ-385/MZ-875 and MZ-383/MZ-876 (Supplementary table S1) for the N- and C-terminal fragments flanking the region to be deleted, respectively. The two PCR products, possessing a 49 bp overlap, were then used as a template for subsequent PCR with the MZ-383/MZ-385 primers, yielding the full-length fragment, which was then cloned into a pETDuet vector. Both proteins were expressed in *E. coli* strain BL21(DE3). Cells were grown in LB broth in the presence of ampicillin at 37 °C. When A600 of the cell culture reached 0.5, the incubation temperature was lowered to 16 °C, 0.1 mM IPTG were added, cells incubated for approx. 16 hours at 16 °C and harvested by centrifugation.

### Protein purification

Harvested cells expressing (His)_6_-tagged WT AfAgo or the dimerization mutant AfAgoΔ were disrupted by sonication in buffer A (20 mM Tris-HCl (pH 8.0 at 25 °C), 500 mM mM NaCl, 5 mM mercaptoethanol) with 2 mM PMSF (phenylmethylsulfonyl fluoride), incubated for 20 min at 50 °C and cell debris was removed by centrifugation at 48,400 x g for 1 hour. The supernatant was loaded onto a HiTrap chelating HP column charged with Ni^2+^ (GE Healthcare) and eluted with a linear gradient (15-500 mM) of imidazole in buffer A. The fractions containing protein were pooled, diluted to 0.2 M of NaCl with a buffer containing 20 mM Tris-HCl (pH 8.0 at 25 °C), 10% glycerol, 5 mM 2-mercaptoethanol and incubated for 1 h at 37 °C with 1 mM EDTA (ethylenediaminetetraacetic acid) and RNase A/T1 (ThermoFisher Scientific) (1:100). Next, the protein solution was centrifuged at 48,400 x g for 30 minutes, the supernatant containing RNA-free AfAgo was loaded onto a HiTrap Heparin HP column (GE Healthcare), and eluted using a 0.2 – 1.0 M NaCl gradient. Finally, the protein was run through the HiLoad 16/600 Superdex 200 column (GE Healthcare) in buffer A and dialyzed against 20 mM Tris-HCl (pH 8.0 at 25 °C), 500 mM NaCl, 50% glycerol.

### Small angle X-ray scattering

Small angle scattering data of WT AfAgo and monomeric mutant were collected at the P12 EMBL beamline on the PETRA III ring of the DESY synchrotron in Hamburg, Germany [36]. Details of data collection and principal structural parameters are presented in Supplementary table S2 and Supplementary figure S1. Protein complexes with DNA (MZ-1289) were transferred to sample buffer (20 mM Tris-HCl (pH 7.5 at 25 °C), 5 mM MgCl_2_, 150 mM NaCl and 2 mM 1,4-dithiothreitol) using Illustra NAP columns (GE Healthcare). Apo AfAgo sample was measured in the same buffer containing 500 mM NaCl.

Dimeric AfAgo complex with MZ-1289 was analyzed by SEC-SAXS with FPLC (Agilent) using Wyatt-MALLS-DLS detection system. The AfAgo+MZ-1289 was concentrated to 175 μM and loaded on the column Superdex 200 Increase 10/300 (GE Healthcare) equilibrated with the sample buffer. Frames collected during the complete SEC run (flow rate 0.5 ml/min, 3000 frames) were analyzed with CHROMIXS [37], frames corresponding to the peak were averaged and processed. *Ab initio* shape determination was carried out by generating 20 independent DAMMIF [38] models using parameterized scattering curves created by GNOM [39] under P2 symmetry restraints. Models were clustered by DAMCLUST [40] and models forming a cluster were averaged by DAMAVER [41] and used as a starting model for an additional run of DAMMIN [42].

SAXS measurements performed with a range of AfAgoΔ concentrations (1-10 mg/ml, both the apo-form and a complex with MZ-1289 DNA), showed significant protein aggregation, particularly pronounced with the apo-AfAgoΔ. The pseudo-chain dummy residues models of the complex generated by GASBOR [43] were superimposed with crystallographic dimers of AfAgo as well as with the monomeric AfAgo-DNA complex using SUPCOMB [44] applying step-wise shift (5 Å) along the principal axis of the model as described in [45].

Comparison of the SAXS data with crystal structures was carried out by CRYSOL [46] (Figure 1). Particle volume and MW estimations were performed using several methods (Supplementary table S3 and references therein).

### DNA fragments

DNA fragments were assembled and prepared as depicted in Supplementary figure S2. All full-length DNA fragments were subsequently purified from an agarose gel using a runVIEW system (Cleaver Scientific, UK), precipitated with sodium acetate/isopropanol, washed with 75% ethanol and resuspended in water.

### AFM sample preparation and imaging

DNA-protein complexes were formed by incubating the DNA fragment (5 nM) with WT AfAgo or AfAgoΔ (concentration in terms of monomer 50 nM) for 5 min at room temperature in the Binding Buffer HEPES (33 mM HEPES (pH 7.8 at 25 °C), 66 mM CH_3_COOK, 5 mM (CH_3_COO)_2_Mg,) in a total volume of 50 μl. Next, the protein-DNA complexes were cross-linked with 2.5% glutaraldehyde for 20 min. Glutaraldehyde was then quenched with an excess of the Tris buffer (33 mM Tris-acetate (pH 7.8 at 25 °C), 66 mM CH_3_COOK, 5 mM (CH_3_COO)_2_Mg,). The resultant reaction solution after 10-fold dilution with Tris buffer was deposited onto modified mica at room temperature as described below.

Freshly cleaved muscovite mica (grade IV, SPI supplies Inc., USA) was incubated in a mixture of 1-(3-aminopropyl)-silatrane (APS) solution for 30 min to prepare functionalized APS-mica, as described previously for the preparation of protein-DNA complexes [47]. 50 μl of DNA-protein complex solution was deposited on APS-mica for 5 min. After incubation the mica surface was immersed into deionized water for 5 min, flushed with excess water and then dried under a flow of nitrogen. The images were acquired in the air with ‘DimensionIcon’ (Bruker, Santa Barbara, CA) microscope system in tapping mode. Probes with nominal spring constants of ~ 5 or 40 N/m were used. Typically, the images were collected at a speed of 0.6 Hz and a resolution of 1024 × 1024 pixels, scan size 2 μm x 2 μm.

### Single-molecule fluorescence microscopy

The overall idea of fluorescence burst data acquisition of single diffusing molecules in alternating laser excitation (ALEX) mode was based on [48]. The principal opto-mechanical layout of the experiment is shown in Supplementary figure S3 and described in Supplementary Methods.

The measurement of single surface-immobilized molecules with the excitation in the total internal reflection mode (TIR) was performed on the same setup exploiting its alternative functionality as described previously [21]. Briefly, the objective was 100× 1.4 Oil Plan Apo VC (Nikon), the fluorescence signal was split by T640lpxr-UF2 dichroic mirror (Chroma) and the different spectral channels were projected on the same EMCCD (DU-897ECS0-UVB, Andor).

### Sample cell preparation for single molecule measurements

FRET bursts measurements were performed in a chambered coverglass well (155411, Nunc Lab-Tek, Thermo Scientific). The reaction volume was 200 μl. The reaction buffer (RB) was 33 mM Tris-acetate (pH 7.9 at 20 °C), 66 mM CH_3_COOK, 5 mM (CH_3_COO)_2_Mg, and 0.1 mg/ml BSA (bovine serum albumin). The DNA concentration was 17-50 pM. Measurements at different protein concentrations were carried out by adding to the reaction small volumes of protein diluted in RB in ‘Protein LoBind’ 1.5 mL tubes (Eppendorf). No oxygen-scavenging or triplet-quenching additives were used.

Measurements of surface-immobilized DNA fragments were performed in a flow cell assembled from a six-channel Sticky-Slide VI 0.4 (Ibidi) and a coverslip functionalized with PEG (polyethylene glycol) derivatives as described in detail in [21]. The flow cell was incubated with 5 μg/ml of Neutravidin (Molecular probes) in RB for 2 min, washed with RB, incubated with 5 pM DNA in RB until the density of the surface-immobilized DNA fragments appeared to be appropriate, and washed with RB. For the measurement, 10 nM solution of AfAgo in RB supplemented with 1% glucose (TCI Europe), 2.5 mM Trolox (Sigma-Aldrich), and 15 U/ml glucose oxidase (Sigma-Aldrich) was injected into the cell. Trolox was treated with UV light for 20 min according to Cordes et al. [49]. Single molecule data analysis was performed as described in Supplementary Methods.

## DECLARATIONS

### Autors’ contributions

AS purified the target proteins. EM performed the SAXS measurements and crystallographic analysis. DR and EG performed smFRET experiments. MJ performed AFM measurements. GS, DR, EM and EG drafted the manuscript. MZ designed and coordinated the study, and critically edited the manuscript. All authors read and approved the final manuscript.

## Acknowledgement

Authors gratefully acknowledge prof. Virginijus Siksnys for fruitful discussions and comments on the manuscript. Authors also acknowledge Audra Ruksenaite for mass spectrometry of AfAgo proteins, Tomas Urbaitis for generation of the dimerization mutant AfAgoΔ and initial experiments, and prof. Gintaras Valincius for support. The synchrotron SAXS data was collected at beamline P12 operated by EMBL Hamburg at the PETRA III storage ring (DESY, Hamburg, Germany). We would like to thank Dr. Melissa Graewert for the assistance in using the beamline.

## Supplementary Information

### SUPPLEMENTARY METHODS

#### Single molecule setup

We used a custom single-molecule fluorescence microscopy setup built on a commercial inverted microscope Nikon Eclipse Ti-U equipped with 60× 1.2 WI Plan Apo VC objective (Nikon) used for the excitation and signal collection, two avalanche photodiode (APD)-based single photon counting modules (Tau-SPAD-50, PicoQuant) and 25 mW 532 and 635 nm diode-pumped solid state and diode lasers (Crystalaser), respectively. The laser excitation was reflected off a dichroic mirror (zt532/635rpc-XT, Chroma), and the fluorescence signal filtered off the excitation light with a quadruple-band interference filter (FF01-446/510/581/703, Semrock) and split into two spectral channels with a dichroic mirror (645dcxr, Chroma). ALEX was implemented by directly TTL-modulating the intensity of the 635 nm laser and synchronously modulating the intensity of the 532 nm laser with a mechanical chopper (MC2000B, Thorlabs). The half period of ALEX was 50 μs. Fluorescence photon arrival times were recorded and ALEX was implemented using an FPGA module (PCIe-7851R, National Instruments) and custom Labview (National Instruments) program.

The excitation was focused 50 μm above the sample chamber glass surface. 532 nm excitation intensity was 30 μW, 635 nm - 20 μW. The size of the confocal pinhole was 75 μm. Each measurement was 10 min long.

#### Single Molecule Data analysis

Fluorescence burst analysis was performed using the freely available FRETBursts software [51]. The initial bursts search parameters were m = 10 photons, and F = 6 times the fluorescence background. The total intensity of a burst from both channels and excitation wavelengths was thresholded to be larger than 40 counts, and this yielded ~3000 bursts from a 10 min measurement. Each burst was calculated a proximity ratio, E, according to E = I_d_^a^/(I_d_^a^ + I_d_^d^), here I_d_^a^ and I_d_^d^ are acceptor and donor intensities upon donor excitation, respectively, and stoichiometry parameter, S, according to S = I_d_/(I_d_ + I_a_^a^), here I_d_ is the total donor and acceptor intensity upon donor excitation, and I_a_^a^ is acceptor intensity upon acceptor excitation. Then we built 2D E-S histograms of bursts. Subsequently, bursts with stoichiometry parameter ranging from 0.2-0.9 were selected to build distributions of the proximity ratio, E, of bursts of DNA molecules labelled with both fluorophores only. E histograms were fit with the sum of two Gaussian functions using unconstrained optimization. Then the ratio of the number of looped and unlooped DNA molecules in the ensemble was calculated as the ratio of the area of the Gaussian of high E with that of low E.

The experiment of surface-immobilized DNA fragments was done by first recording a short movie with 635 nm excitation to obtain a reference for fluorescent spot identification since the acceptor channel exhibits significantly less fluorescence background than the donor channel. Then a longer actual movie was recorded with the 532 nm excitation. The analysis of the two-spectral channel fluorescence movies was performed using custom software written in Matlab. Briefly, to identify the fluorescent spots, the first 20 frames of the reference and the actual fluorescence movies were averaged, the obtained average images were filtered with the 2D low-pass Gaussian filter 5 pixels large and with the standard deviation of 1 pixel and subtracted the same image filtered with the averaging filter 7 pixels large. The resulting acceptor channel reference image was thresholded with 20 and the donor channel actual image - with 40 counts/pixel. The obtained images were binarized for particle identification. Particles in both binary images were identified and filtered according to the following criteria: 5×5 pixel ROIs (regions of interest) centered on particles’ centers of mass had to non-overlap, particle area had to be within 5-100 pixels range, particle eccentricity not larger than 0.8. The coordinates of a particle in the donor channel corresponding to a particle identified in the acceptor channel of the reference movie were calculated using the spatial transformation structure calculated from an image of surface-immobilized 200 nm fluorescent polystyrene beads (F8806, Invitrogen). For trace extraction were considered only those particles in the actual movie whose donor coordinates coincided with the transformed coordinates of the acceptor particles in the reference movie within 1.5 pixels. The donor and acceptor intensity traces were extracted using aperture photometry [52] with the background calculated as an average intensity from a 1 pixel-wide annulus around particle’s ROI. The proximity ratio, E, was calculated according to the same formula as for the fluorescence bursts.

**Supplementary figure S1.**
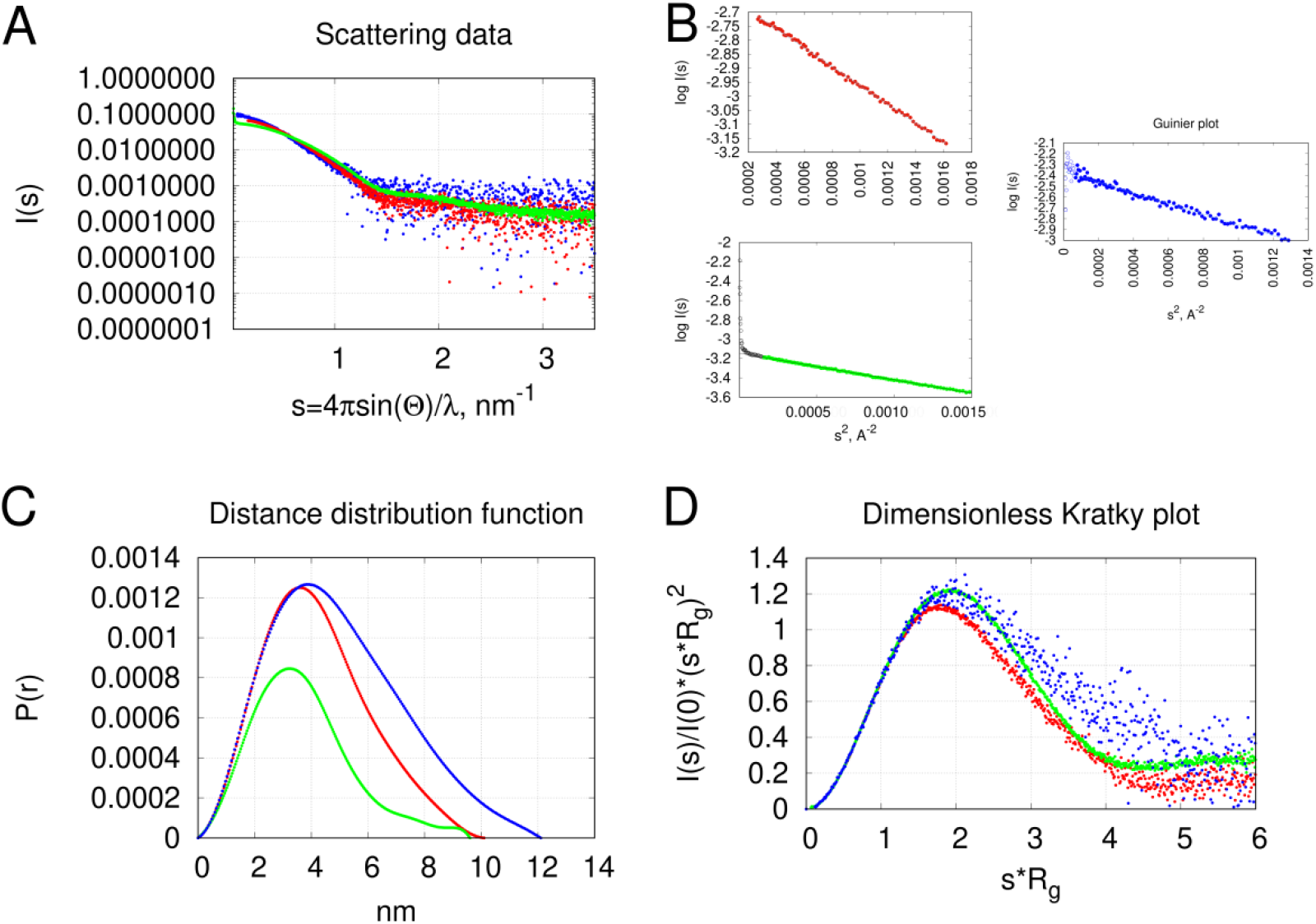
SAXS data of AfAgo+MZ-1289 (red curves), monomeric mutant AfAgoΔ+MZ-1289 (green curves) complexes and apo AfAgo (blue curves). (A), Scattering curves. (B), Guinier plots log I(s) vs. s^2^ of the data at small s values. (C), Pair distance distribution functions. (D), Dimensionless Kratky representation of scattering data (I(s)/I(0)*(s*Rg)^2^ vs. s*Rg). All curves have similar shape typical for folded proteins [53].

**Supplementary figure S2.**
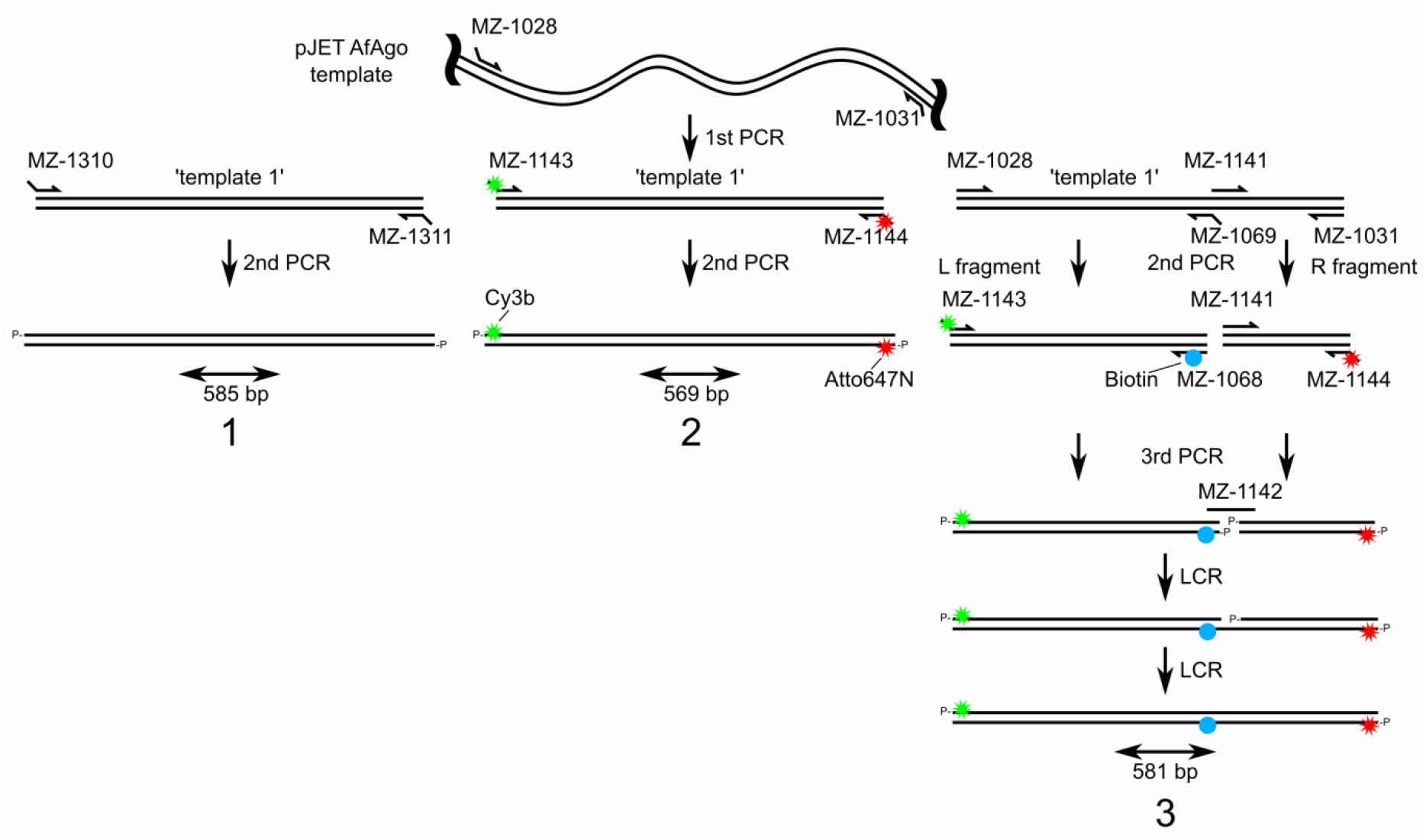
Synthesis scheme for the DNA fragments. First, a DNA fragment was amplified from a pJET plasmid template containing an AfAgo gene fragment using oligonucleotides MZ-1028 and MZ-1031. The PCR product was then used as a template (dubbed ‘template 1’) in subsequent reactions. Fragment ‘1’ used for AFM studies was made by PCR from ‘template 1’, using oligonucleotides MZ-1310 and MZ-1311, which were treated with T4 polynucleotide kinase (PNK) prior to amplification, to yield a 585 bp fragment. Fragment ‘2’ was amplified from ‘template 1’ with oligonucleotides MZ-1143 and MZ-1144, bearing Cy3B (green star) and Atto647N (red star) dyes, respectively, on the third base from the 5’-end, yielding 569 bp DNA. Fragment ‘3’ was synthesised in two steps. Firstly, respective fragments flanking the biotinylation site (dubbed ‘L fragment’ and ‘R fragment’) were amplified by PCR from ‘template 1’, using primer pairs MZ-1028 and MZ-1069 for the ‘L fragment’, and MZ-1031 and MZ-1141 for the ‘R fragment’. Secondly, each of the two fragments were used as templates for subsequent PCRs. ‘L fragment’ was amplified using MZ-1143 and MZ-1068, the latter bearing the biotin (blue circle) on 22 b from its 5’-end. ‘R fragment’ was amplified using primers MZ-1141 and MZ-1144. The two fragments were then purified using a GeneJET PCR purification kit (ThermoFisher Scientific), and treated with PNK while mixed in equal amounts to a total concentration of 6 nM. The phosphorylation mix was subsequently ligated by Ampligase^®^ (Epicentre, USA) at 3 nM total DNA and 30 nM bridging oligonucleotide MZ-1142 according to Chandran, 2017 [54]. All full-length DNA fragments were subsequently purified from an agarose gel using a runVIEW system (Cleaver Scientific, UK), precipitated with sodium acetate/isopropanol, washed with 75% ethanol and resuspended in water.

**Supplementary figure S3.**
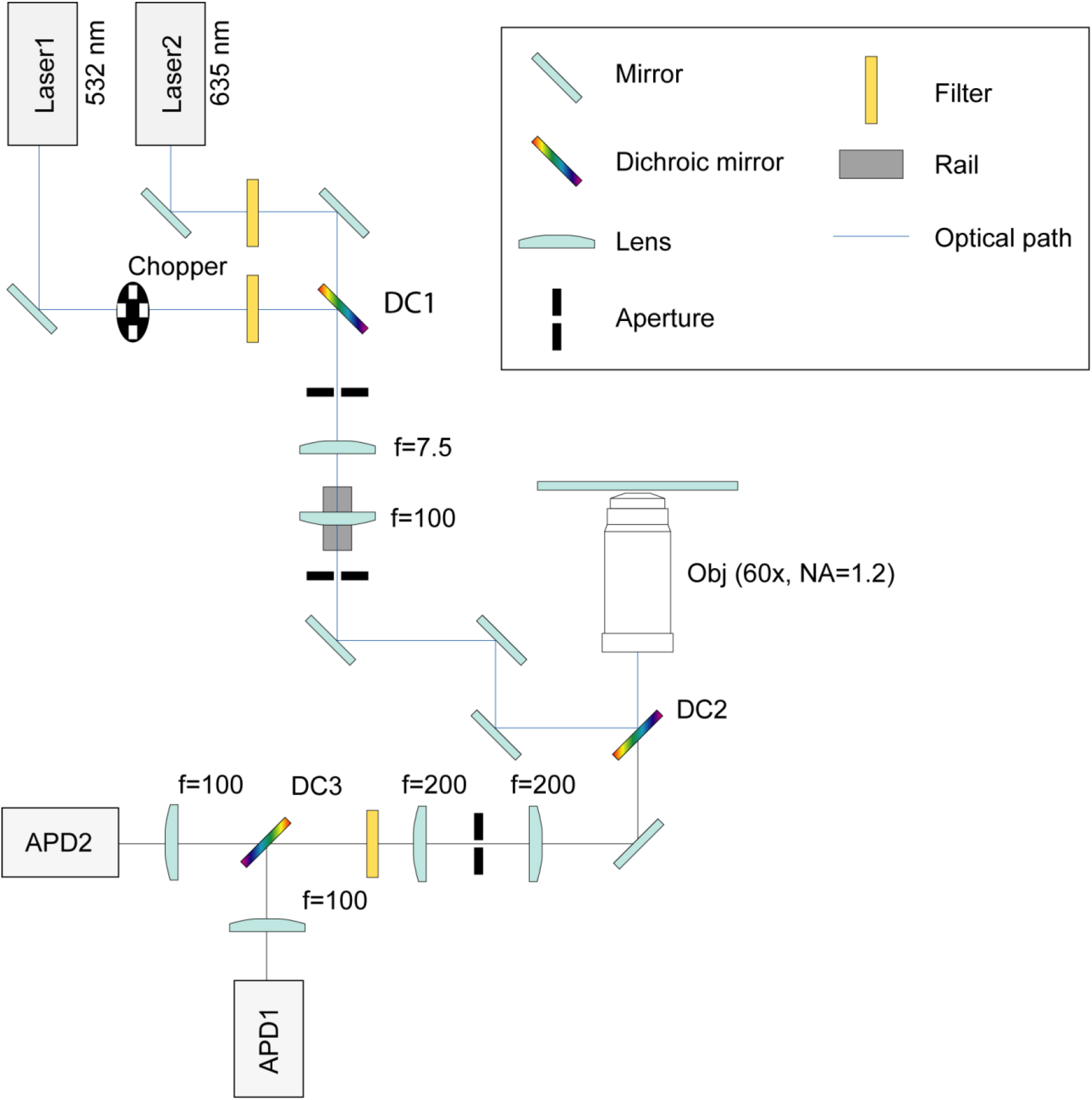
Optical scheme of custom single-molecule fluorescence microscopy setup used to record fluorescence bursts of single diffusing molecules in this study. APD – avalanche photodiode; f – focal distance; NA – numerical aperture.

**Supplementary figure S4.**
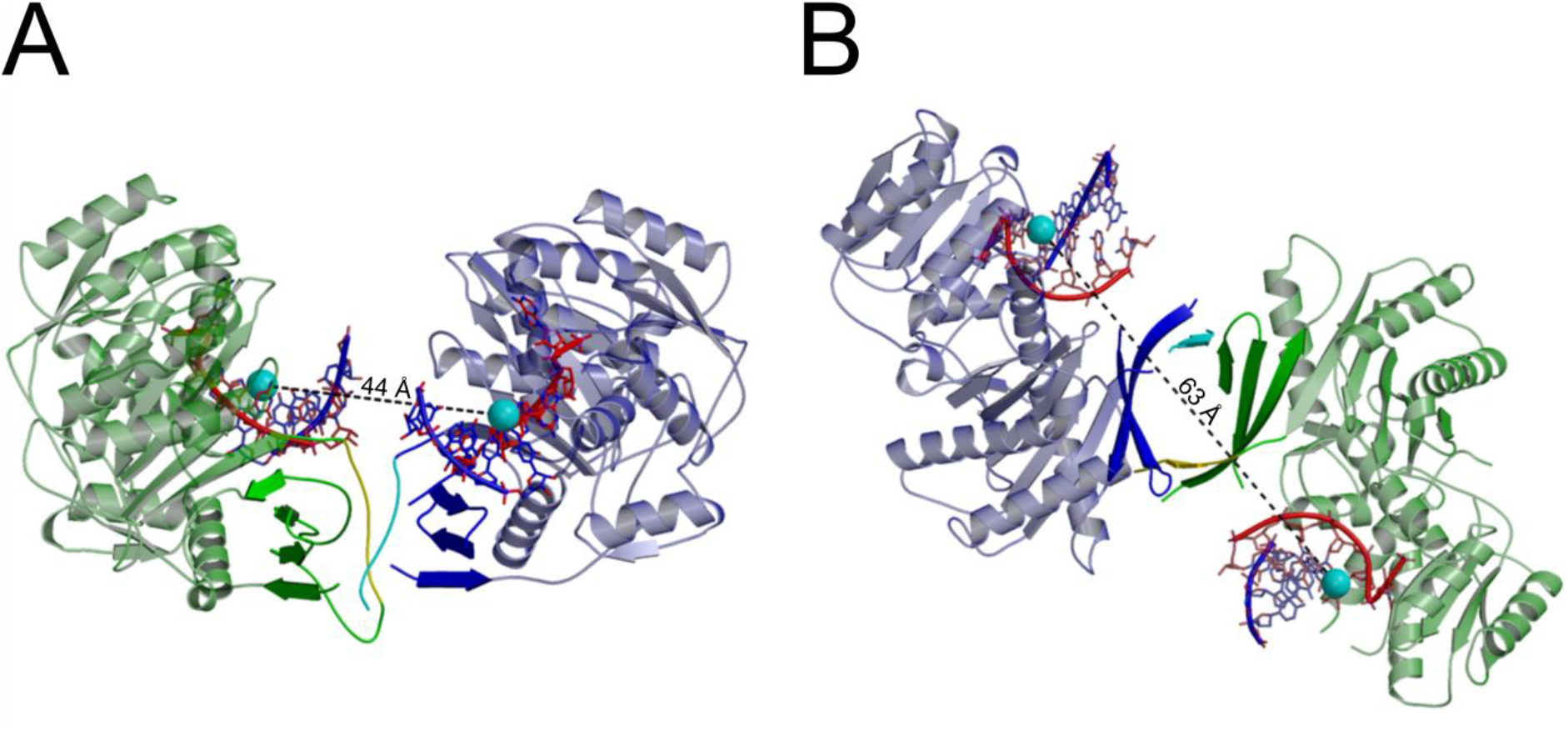
Expected positions of fluorescent labels upon formation of the looped complex. The figure is based on PDB ID 1ytu (A, ‘closed’), 2w42 (B, ‘open’), spheres mark fluorophore attachment sites. Protein monomers are coloured green and blue, DNA guide and target strands are red and blue respectively.

**Supplementary figure S5.**
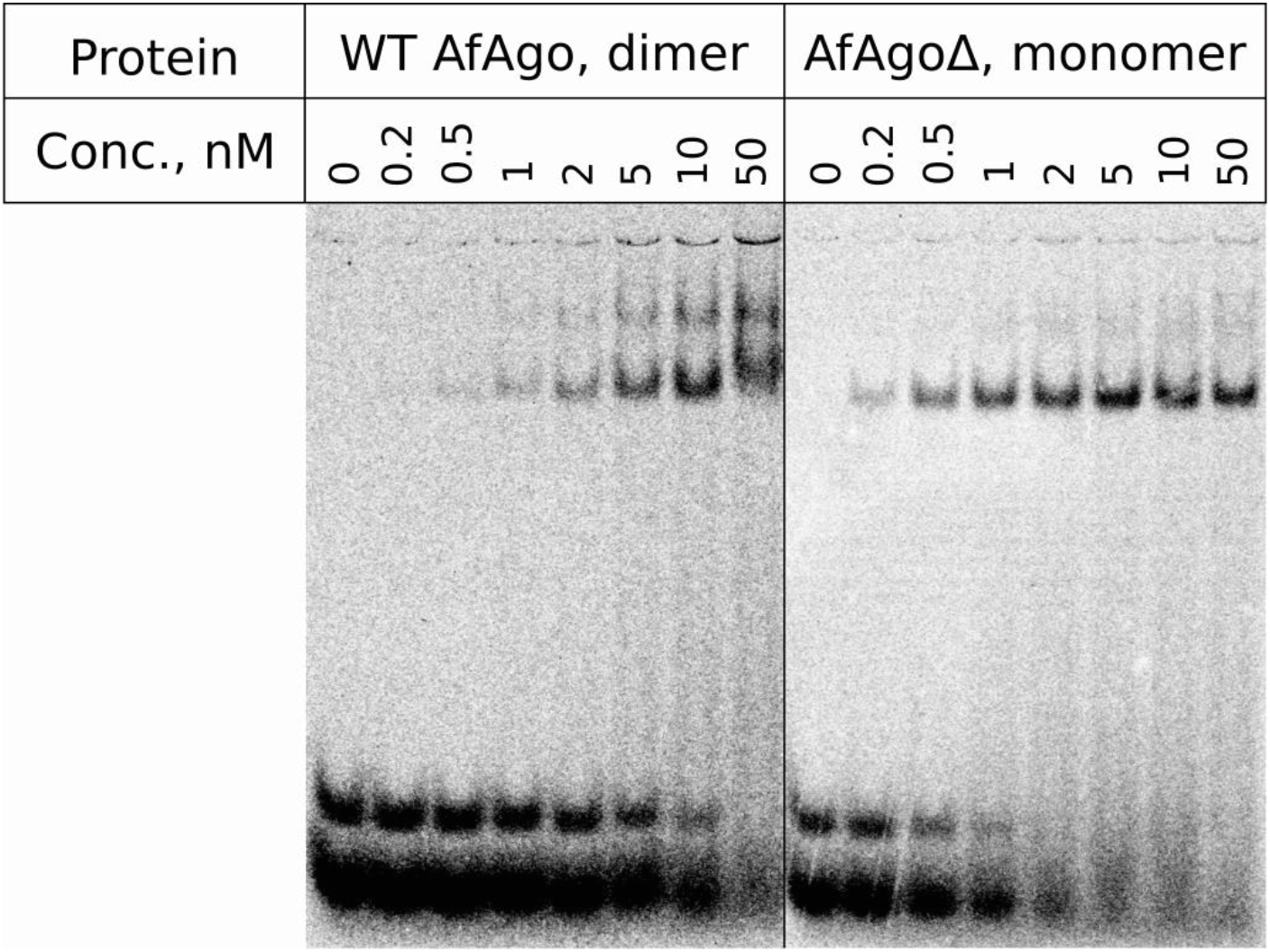
DNA binding by AfAgo. DNA binding was verified using electrophoretic mobility shift assay. Self-complementary 5’P^32^ DNA MZ-952 was used as a substrate. Final concentration of DNA duplex in binding reaction was 1 nM, final protein concentrations are shown above each lane. Binding buffer was 40 mM Tris-acetate (pH 8.4 at 25 °C) with 1 mM EDTA (TAE, B49, Thermo Scientific), supplemented with 5 mM MgCl_2_, 0.1 mg/ml BSA, 1 mM DTT, 10% glycerol. Running buffer – TAE (Thermo Scientific) supplemented with 5 mM MgCl_2_.

**Supplementary figure S6.**
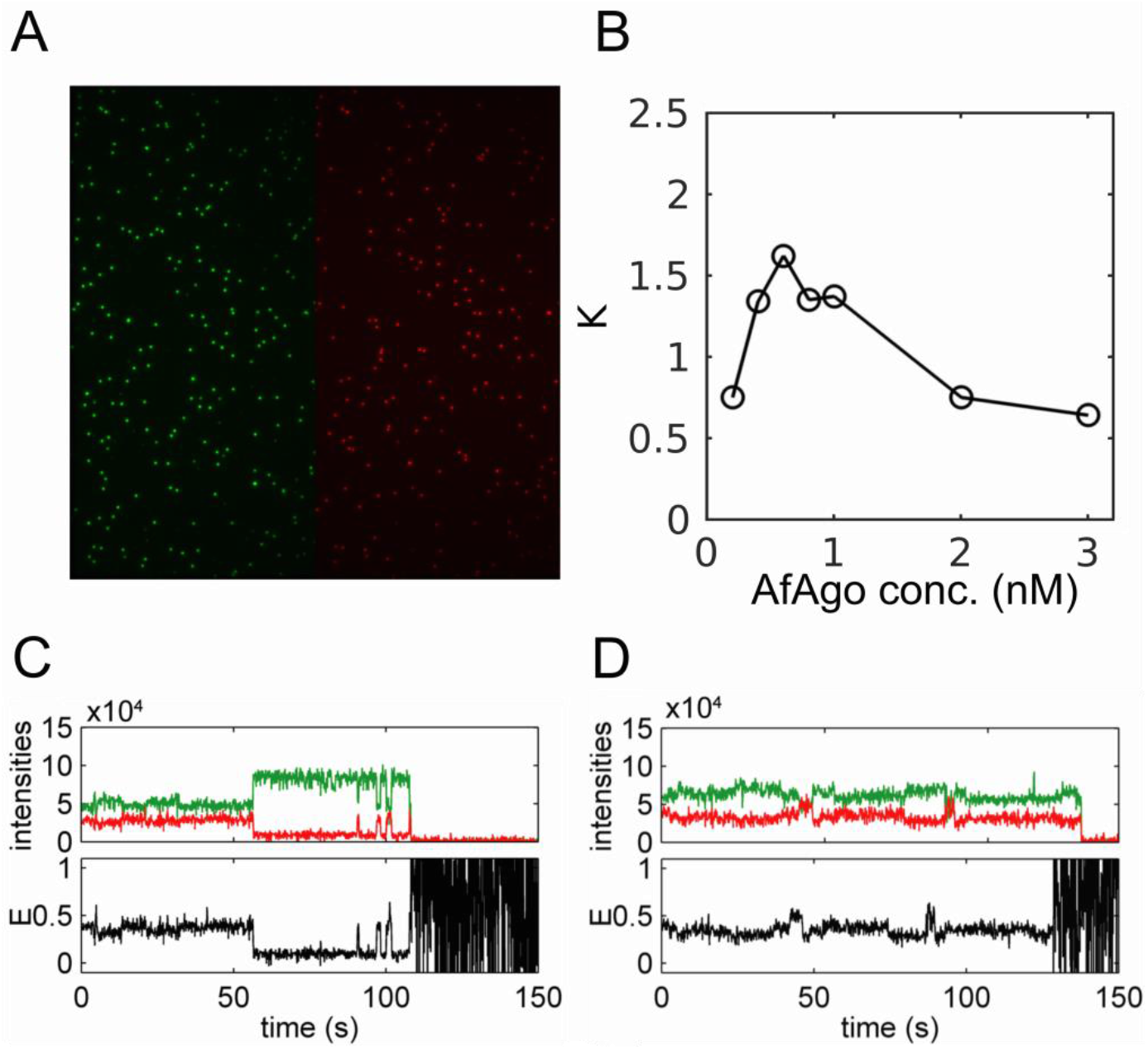
Single molecule experiments. (A) A fluorescence image of surface-immobilized DNA fragments. It is an average of 20 frames in a fluorescence movie. The left part (green) is the donor image upon donor excitation, and the right part (red) is the acceptor image upon acceptor excitation. (B) The dependence of the ratio, K, of the number of looped and unlooped DNA molecules depending on the concentration of the AfAgo for the biotinylated DNA fragment in solution. (C, D) Examples of different dynamics of DNA looping by AfAgo in TIRF experiments.

**Supplementary table S1.**
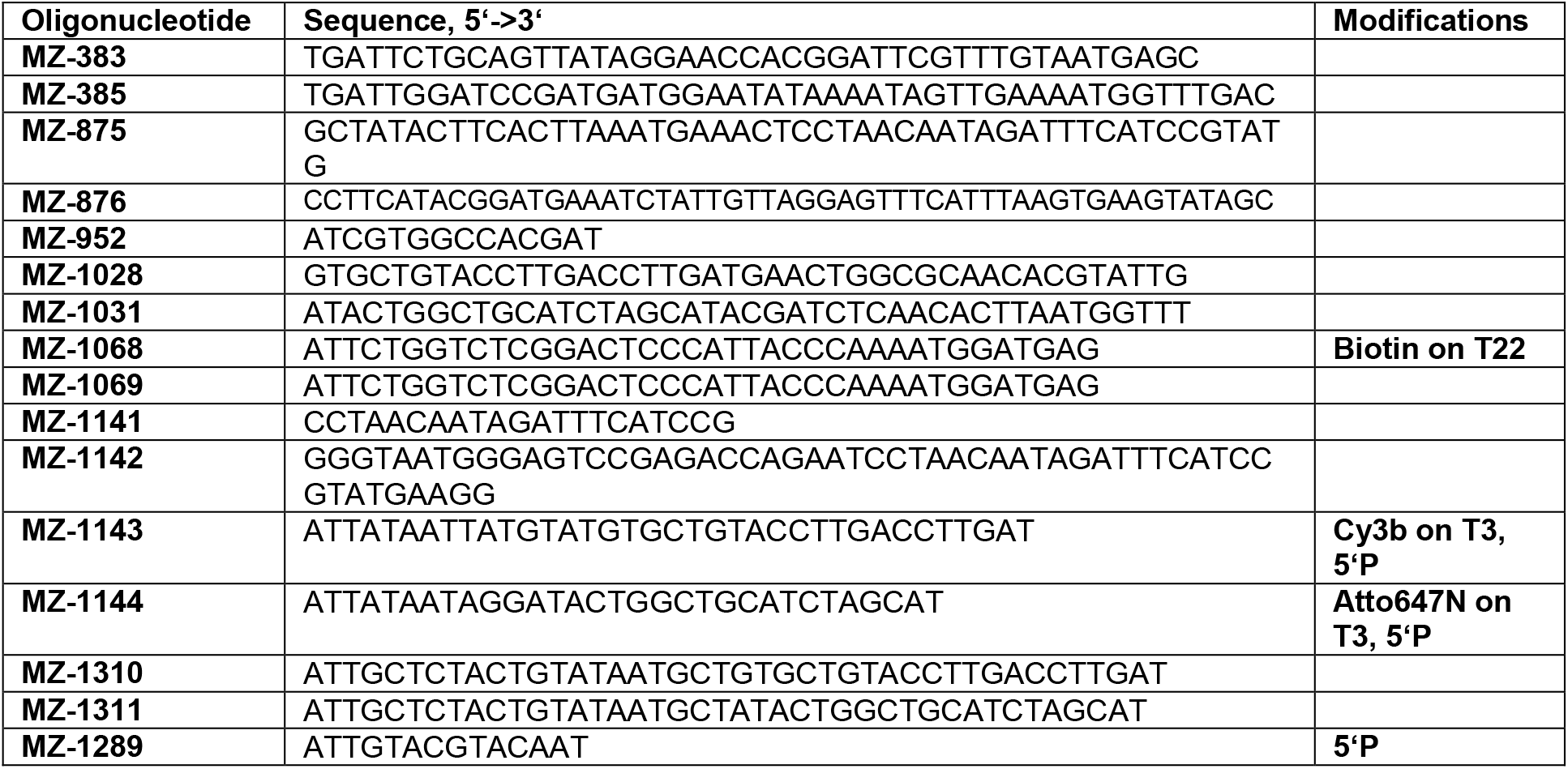
List of oligonucleotides used in this study.

**Supplementary table S2.** SAXS data collection and main structural parameters

|  |  |  |  |
| --- | --- | --- | --- |
| Instrument, Detector | P12, pilatus6m |  | P12, pilatus2m |
| Detector-to-sample distance, m | 3.0 |  | 3.0 |
| Wavelength, nm | 0.123981 |  | 0.124 |
| Measured s range, nm<sup>-1</sup> | 0.0224526-7.3176000 |  | 0.02492870-5.064020 |
| Number of buffer exposure frames averaged (measured) / frame exposure time | 101 (101) / 0.995 sec | 76 (80) / 0.195 sec | 40 (40) / 0.045 |
| Number of sample exposure frames averaged (measured) / frame exposure time | 24 (24) / 0.995 sec | 30 (40) / 0.195 sec | 20 (20) / 0.045 |
| Capillary temperature/ Sample changer temperature | 20 °C / Room temperature | 20 °C / 10 °C | 20 °C / 10 °C |
| Data reduction and on-line characterization | radaver (r11095), databsolute v0.1 (r11095) |  | radaver (v. 9729) |
| Structural parameters |  |  |  |
| Sample | WT AfAgo+MZ-1289, SEC peak | AfAgoΔ+MZ-1289, 4 mg/ml | Apo AfAgo, 0.13 mg/ml |
| Guinier points (AUTORG) | 1-87 | 39-132 | 20-114 |
| s range, nm<sup>-1</sup> (points) used in GNOM | 0.0640-3.3457 (1-1200) | 0.1860-3.3457 (60-1200) | 0.0249-2.8342 (20-1000) |
| Rg, nm (AUTORG/ GNOM) | 3.18 ± 0.016/<br>3.233 ± 0.005202 | 2.84 ± 0.03/<br>2.879 ± 0.002440 | 3.83 ± 0.12 /<br>3.837 ± 0.02216 |
| I(0) (AUTORG/ GNOM) | 0.0725 ± 0.00011/<br>0.07301 ± 0.00008771 | 0.0428 ± 3.7e-05/<br>0.04289 ± 0.00002499 | 0.0935 ± 0.00043 /<br>0.09310 ± 0.0005914 |
| Dmax, nm (DATCLASS/ SHANUM/ GNOM) | 11.3/ 10.2/ 10.1 | 10.9/ 10.5/ 9.6 | 13.9/ 11.5/ 12.1 |
| Porod volume, nm<sup>3</sup> (DATPOROD) | 158.03 | 108.67 | 174.1 |
| SASBDB ID | SASDH39 | SASDH49 | - |

**Supplementary table S3.** Molecular mass determination from SAXS data using various methods. All molecular masses are given in kDa

| Sample |  |  | WT AfAgo+MZ-1289 | AfAgoΔ+MZ-1289 | Apo AfAgo |
| --- | --- | --- | --- | --- | --- |
| Expected MW (protein + DNA), kDa |  |  | 110.5 | 56.6 | 101.7 |
| Method | Reference | Software | MWcalc |  |  |
| Absolute scale | [55] | PRIMUS 2.8.4<br>(r10552) | 99.7 | 55.4 | 130.2 |
| Qp |  |  | 102.7 | 58.5 | 128.8 |
| Bayes |  |  | 94.2 | 56.9 | 124.5 |
| Size&Shape |  |  | 100.0 | 67.9 | 133.1 |
| Porod volume/1.6 | [40] | DATPOROD,<br>ATSAS 2.8.4<br>(r10552) | 98.8 | 67.9 | 108.8 |
| SAXSMoW | [56] | SAXSMoW<br>v2.1<br>http://saxs.ifsc.usp.br/ | 106.9 (integrated to<br>$I_0/I(q_{max})=10^{2.25}$) | 67.4<br>(integrated to<br>$I_0/I(q_{max})=10^{2.25}$) | 125.0<br>(integrated to 8/Rg) |
| SEC MW |  | CHROMIXS<br>ATSAS 2.8.4<br>(r10552) | 103.8 | n.a. | n.a. |

**Supplementary table S4.**
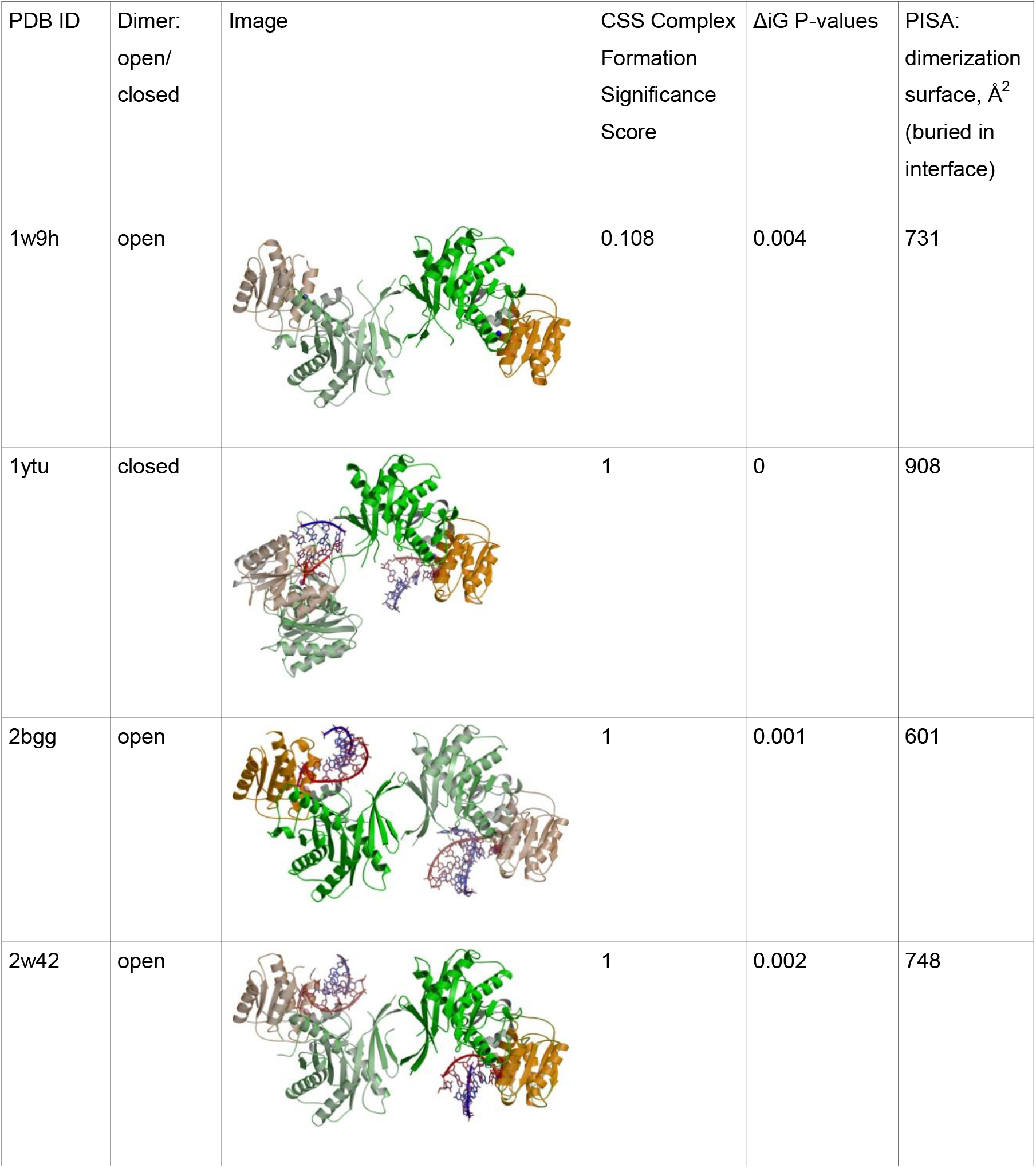
AfAgo dimerization interfaces as analyzed by PISA (PDBe PISA v1.52 [20/10/2014])

| PDB ID | Dimer:<br>open/<br>closed | Image | CSS Complex<br>Formation<br>Significance<br>Score | ΔiG P-values | PISA:<br>dimerization<br>surface, Å<sup>2</sup><br>(buried in<br>interface) |
| --- | --- | --- | --- | --- | --- |
| 1w9h | open |  | 0.108 | 0.004 | 731 |
| 1ytu | closed |  | 1 | 0 | 908 |
| 2bgg | open |  | 1 | 0.001 | 601 |
| 2w42 | open |  | 1 | 0.002 | 748 |

